# Chagas disease serological test performance in United States blood donor specimens

**DOI:** 10.1101/730754

**Authors:** Jeffrey D. Whitman, Christina A. Bulman, Emma L. Gunderson, Amanda M. Irish, Rebecca L. Townsend, Susan L. Stramer, Judy A. Sakanari, Caryn Bern

## Abstract

**Background:** Chagas disease affects an estimated 300,000 individuals in the US. Diagnosis in the chronic phase requires positive results by two different IgG serological tests. Three ELISAs (Hemagen, Ortho, Wiener) and one rapid test (InBios) are FDA-cleared, but comparative data in US populations are sparse.

**Methods:** We evaluated 500 seropositive and 300 seronegative blood donor plasma samples. Country of birth was known for 255 seropositive specimens and grouped into regions: Mexico (n=94), Central America (n=88) and South America (n=73). Specimens were tested by the four FDA-cleared IgG serological assays. Test performance was evaluated by two comparators and latent class analysis.

**Results:** InBios had the highest sensitivity (97.4-99.3%), but lowest specificity (87.5-92.3%). Hemagen had the lowest sensitivity (88.0-92.0%), but high specificity (99.0-100.0%). Sensitivity was intermediate for Ortho (92.4-96.5%) and Wiener (94.0-97.1%); both had high specificity (98.8-100.0% and 96.7-99.3%, respectively). Antibody reactivity and clinical sensitivity was lowest in donors from Mexico, intermediate in those from Central America and highest in those from South America.

**Conclusions:** Our findings provide an initial evidence base to improve laboratory diagnosis of Chagas disease in the US. The best current testing algorithm would employ a high sensitivity screening test followed by a high specificity confirmatory test.

## Introduction

Chagas disease is the most important tropical disease in the Americas. The attributable disease burden in the region, based on disability-adjusted life years lost, is nearly 8 times greater than that due to malaria and 20% higher than that for dengue (1). An estimated 6 million people are currently infected with *Trypanosoma cruzi*, predominantly in Mexico, Central and South America (2). Chronic infection persists lifelong in the absence of treatment, with tissue tropism for cardiac myocytes and the enteric nervous system (3-6). Over time, 20-30% of infected individuals develop cardiac or gastrointestinal disease. The United States (US) contains widespread enzootic transmission cycles involving wildlife and sylvatic triatomine vectors, but autochthonous *T. cruzi* transmission to humans appears to be very rare (7, 8). Locally acquired infections are greatly outnumbered by the estimated 300,000 infected immigrants from Latin America residing in the US (9, 10). The US Food and Drug Administration (FDA) approved benznidazole, the first-line Chagas disease treatment, in 2017, increasing the need for reliable diagnostic testing for both individual and public health needs in the US (11).

In the chronic phase, confirmed diagnosis requires positive results by two serological tests for IgG antibodies to *T. cruzi*, preferably based on different antigens (12). Currently, four serological assays are cleared by FDA for diagnostic use, Ortho *T. cruzi* ELISA (Ortho Clinical Diagnostics, Raritan, NJ), Hemagen Chagas’ Kit ELISA (Hemagen Diagnostics, Inc, Columbia, MD), Wiener Chagatest Recombinante v.3.0 ELISA (Wiener Laboratories, Rosario, Argentina), and InBios Chagas Detect Plus rapid test (InBios International, Inc, Seattle, WA) (13). All four assays report high sensitivity and specificity in their FDA 510(k) clearance applications (reported percent sensitivity/specificity: Ortho 98.9/99.99; Hemagen 100/98.7; Wiener 99.3/98.7; InBios 95-100/87-98). However, comparative performance data are lacking for at-risk populations in the US, as well as those in Mexico and Central America, the predominant regions of origin of US immigrants (14). Emerging evidence suggests variation in test sensitivity by geographic location and a high rate of discordance between serological tests, particularly in Mexico (15-18). Comprehensive studies are needed to provide the basis for development of reliable testing algorithms. In this study, we compared the performance of the four FDA-cleared serological tests in specimens from US blood donors to provide the first systematic evidence to improve laboratory diagnosis of Chagas disease in the US.

## METHODS

### Ethical Approval

This study was approved by ARC institutional review board and deemed exempt from review by the Human Research Protection Program at UCSF.

### Sample Selection and Preparation

We evaluated archived plasma samples from 800 blood donations collected by the American Red Cross (ARC) between September 2006 and June 2018. Specimen selection was based on confirmed *T. cruzi* infection status in ARC blood donation (BD) testing algorithms at the time of blood donation (8). ARC provided a list of 1091 seropositive specimens, defined by repeat reactive results by an FDA-licensed screening test (Ortho ELISA or Abbott (Abbott Laboratories, Abbott Park, IL) PRISM) followed by confirmed-positive results by a supplemental test (radioimmunoprecipitation assay [RIPA] (performed by Quest Diagnostics (Chantilly, VA)) or Abbott Enzyme Strip Assay [ESA]) (8). We prioritized selection of BD-positive specimens with country of birth data; the remainder of the 500 BD-positive specimens were selected at random. A random sample of 300 specimens was compiled from a list of 3938 seronegative blood donations, frequency-matched by region of donation to the BD-positive specimen set. No country of birth data were available for seronegative specimens.

Donated plasma units from each donation were frozen at −20°C within 24 hours of collection. Retrieved plasma units from ARC collections used for research purposes were thawed in a temperature-controlled water bath, aliquoted into multiple tubes and refrozen. Aliquots tested by Hemagen, Wiener and InBios at UCSF were thawed and refrozen only once. For the current analysis, the Ortho ELISA was re-run on all 800 specimens in 2019. Aliquots used for current Ortho testing were thawed and refrozen twice.

Ortho ELISA testing for this study was conducted at Innovative Blood Resources, Minneapolis, MN, using the fully automated Ortho Summit System (19). The Ortho ELISA has FDA approval for blood donation screening and clearance for diagnostic purposes, but is not yet marketed for the latter use. For the Ortho ELISA, signal-to-cutoff (S/CO) ratios of 1.00 or greater are considered reactive; in the blood donation screening algorithm, all reactive units are retested two more times. A blood donation is considered repeat reactive if at least 2 of 3 sample results have an S/CO greater than 1.00.

Testing by Hemagen ELISA, Wiener ELISA, and InBios rapid tests was conducted at the University of California, San Francisco (UCSF), San Francisco, CA. Plasma samples were thawed at 4°C and spun at 2300 relative centrifugal force for 10 minutes to pellet any precipitate. Samples were aliquoted to randomly assigned positions in 96 deep-well plates to blind readers performing the InBios rapid test. Plasma aliquots of 10µL (Hemagen, Wiener) or 5µL (InBios) were tested and interpreted in accordance with package inserts using the kit reagents, ELx405 Select Microplate Washer (BioTek, Winooski, VT), and SpectraMax Plus 384 Microplate Reader (Molecular Devices, San Jose, CA). The InBios package insert defines a positive result as any visible test line. To quantify this assay a set of 7 quality control samples was used to construct a semi-quantitative scale from 0 (negative) to 6 (strong positive) (Figure S1). InBios test results were scored by two independent readers blinded to other assay results. The only deviation from package insert protocols was the use of plasma for Hemagen tests; the manufacturer recommends use of serum only.

### Data Analysis

We conducted three analyses to assess diagnostic test performance. Two analyses compared assay results to different reference standards: classification in prior BD testing and a consensus classification based on positive results by two or more diagnostic assays in the current study. BD was defined by the blood donation testing algorithms described above (8). For InBios testing, reader 1 scores were used for performance calculations; reader 2 scores were used to calculate inter-reader agreement statistics. The Hemagen and Wiener kits both include an indeterminate zone; results that fell in this zone were included as positive in the performance analyses, because they would necessitate confirmatory testing in real-world scenarios. This definition may overestimate the sensitivity and/or specificity of these two tests (depending on whether the grey zone results predominantly correspond to seropositive or seronegative specimens). Exact binomial 95% confidence intervals (CI) were calculated for each of the performance parameters. Analyses were conducted in SAS 9.4 and R version 3.5.2.

The third performance assessment consisted of a latent class analysis (LCA). LCA comprises a group of mathematical modeling techniques developed to evaluate diagnostic tests in the absence of a true gold standard (20-23). We assumed two latent classes and conditional independence of test outcomes. We used bootstrapping to generate multiple samples from the dataset and then applied an expectation-maximization (EM) algorithm to estimate sensitivity and specificity for each test. The distributions of the bootstrapped samples were used to generate 95% CIs. We tested the robustness of the two-class assumption by comparing fit between models assuming two versus three latent classes, using the Akaike information criterion (AIC) and Bayesian information criterion (BIC). The latent class analysis was conducted in R version and RStudio version 1.1.463 using the BayesLCA package (24).

## RESULTS

California and the southeastern states accounted for nearly three-quarters of the blood donations included in the study (Table 1). BD-positive specimens were significantly more likely than BD-negative ones to be from donors who identified themselves as Hispanic. Among 282 BD-positive donors with country of birth data, 33% were from Mexico, 31% from Central America and 26% from South America. Approximately 10% of donors with country of birth data were born in the US, but the source of their infections was likely a mixture (congenital, travel or locally acquired); this group of donations was not included in analyses by birth country.

**Table 1.** Characteristics of donors whose specimens were used in the evaluation.

|  | Blood donor status |  |
| --- | --- | --- |
|  | Positive N (%) | Negative N (%) |
| Region of blood donation<sup>1</sup> |  |  |
| California | 195 (39.0) | 115 (38.3) |
| Other western states | 50 (10.0) | 31 (10.3) |
| Southeast | 170 (34.0) | 103 (34.3) |
| Midwest | 33 (6.6) | 19 (6.3) |
| Northeast | 52 (10.4) | 32 (10.7) |
| Sex |  |  |
| Male | 240 (48.0) | 127 (42.3) |
| Female | 260 (52.0) | 173 (57.7) |
| Ethnicity |  |  |
| Hispanic<sup>2</sup> | 204 (40.8) | 37 (12.3) |
| Caucasian | 17 (7.6) | 223 (74.3) |
| Other | 1 (0.5) | 35 (11.7) |
| No data | 278 (55.6) | 5 (1.7) |
| Region of birth<sup>3</sup> |  |  |
| Mexico | 94 (33.3) |  |
| Central America | 88 (31.2) |  |
| South America | 73 (25.9) |  |
| USA | 27 (9.6) |  |

<sup>1</sup>Seronegative specimens frequency-matched to seropositive specimens by donation region.
<sup>2</sup>Positive blood donors significantly more likely to report Hispanic ethnicity ( $p < 0.0001$ ).
<sup>3</sup>Data available for 282 blood donors identified as seropositive in blood donation testing; no data for 218 seropositive and 300 seronegative specimens.

The three analyses (BD-status, consensus, and LCA) yielded similar results, with overlapping 95% CIs for each parameter across analyses of the same test (Table 2). The highest sensitivity estimates resulted from the LCA and the lowest from the BD comparison; the reverse trend was seen for specificity. The 2-class LCA showed better fit than a 3-class analysis both by AIC (−2059.089 vs −2027.283) and BIC (−2101.25 vs −2092.867).

**Table 2.**
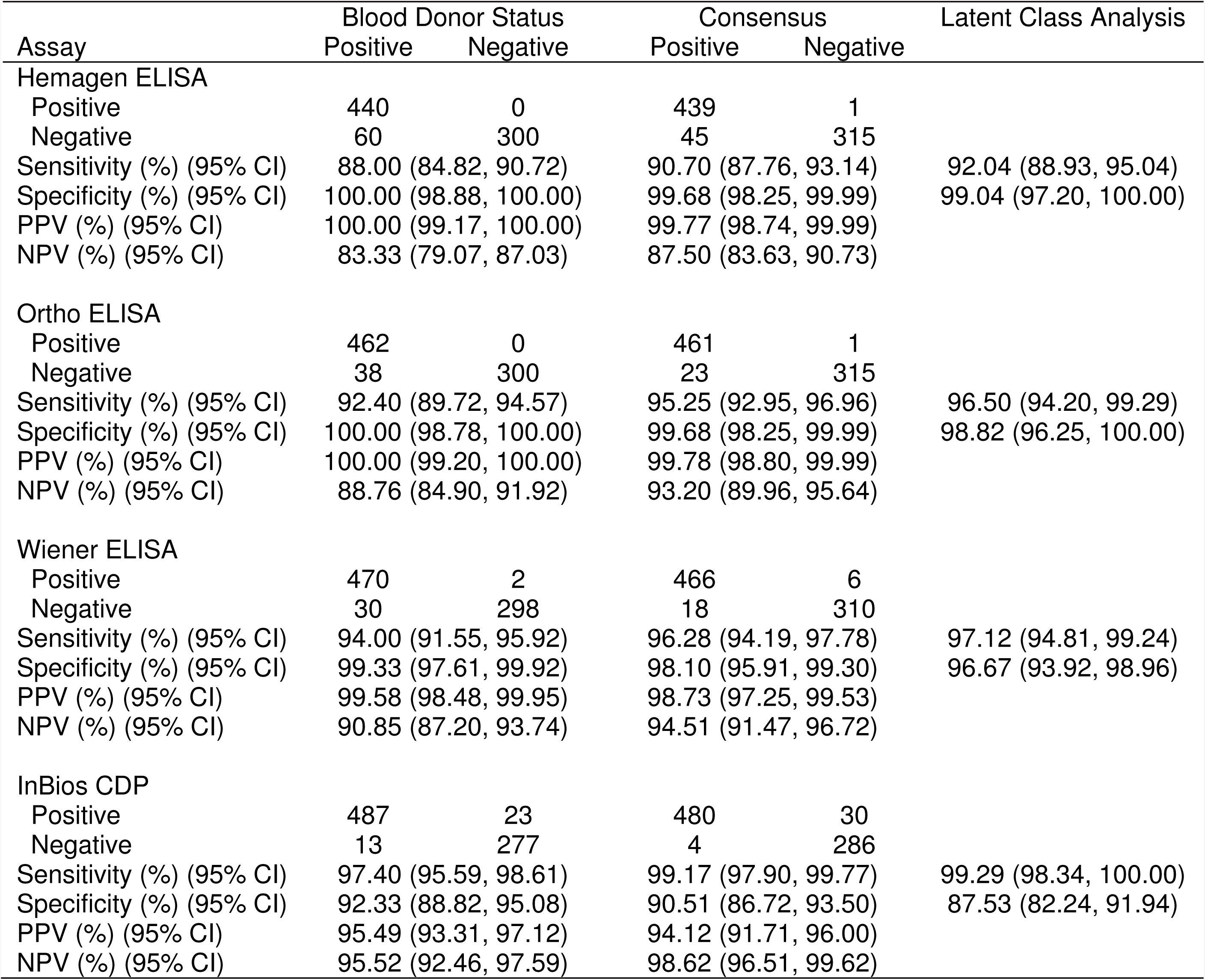
Performance of FDA-Cleared Chagas disease IgG serological ssays compared to original blood donor status, consensus among current tests, and latent class analysis.

In all three analyses, InBios CDP had the highest sensitivity (97 to 99%), but the lowest specificity (88 to 92%). Reader agreement on InBios scores was high (weighted kappa=0.9315 (95% CI 0.9209, 0.9420). Agreement on determination of positive (score 1-6) versus negative (score 0) was more than 99% (795/800 (99.4%); kappa=0.9865 [95% CI 0.9746, 0.9983]. There were only five discordant results: two specimens positive by reader 1 and negative by reader 2, three specimens with the converse. The majority of apparent false-positive InBios results had intensity scores of 1 (87% for BD, 83% for consensus analysis). Hemagen displayed the lowest sensitivity (88 to 92%) but high specificity (99 to 100%). Eleven specimens had Hemagen readings in the indeterminate zone; all were BD-positive. Sensitivity for the Wiener ELISA ranged from 94 to 97%, with specificity 97 to 99%. Six specimens had indeterminate results by Wiener, four BD-positive and two BD-negative. Of the 500 specimens classified as confirmed positive in BD testing, those with negative results by current assays (apparent false negatives) had significantly lower median Ortho S/CO values in prior BD testing compared to those with positive results in current testing (apparent true positives) (Figure S2).

Ortho ELISA sensitivity ranged from 92 to 97% in the current analysis, with specificity of 99 to 100%. Of 500 BD-positive specimens, 489 had positive Ortho results in BD testing; 11 specimens were positive by Abbott PRISM and a supplemental test (RIPA and/or Abbott ESA) but negative by Ortho in BD testing. Four of the 11 previously Ortho-negative specimens had positive results in the current Ortho testing, but 31 previously Ortho-positive specimens had negative results. Current Ortho S/CO values were a median 15.9% lower than in BD testing (p<0.001). Specimens corresponding to earlier collected donations showed a smaller decline in S/CO values than more recent ones (Y = 0.007334*X - 1.534; R^2^ = 0.05758; p< 0.001 in linear regression analysis of percent decline in S/CO vs specimen age in months).

Finally, we stratified results by region of birth to explore geographic variation in test sensitivity (Table 3). Compared to BD or consensus status, sensitivity for Ortho, Wiener and Hemagen tended to be lowest in specimens from those born in Mexico and highest in those from South America, with Central American specimens showing intermediate results. Analyses of the antibody reactivity were consistent with these results, with the lowest reactivity in Mexico (Figure 1).

**Table 3.** Sensitivity of *T. cruzi* IgG serological tests by blood donor region of birth

| Test | Blood donor status |  |  | Consensus (at least 2 current tests positive) |  |  |
| --- | --- | --- | --- | --- | --- | --- |
|  | Mexico | Central America<sup>1</sup> | South America<sup>2</sup> | Mexico | Central America<sup>1</sup> | South America<sup>2</sup> |
| Hemagen | 82.98<br>(74.13, 89.24) | 88.64<br>(80.33, 93.71) | 93.15<br>(84.95, 97.04) | 86.67<br>(78.13, 92.21) | 89.66<br>(89.66, 94.46) | 93.15<br>(84.95, 97.04) |
| Ortho | 85.11<br>(76.54, 90.92) | 95.45<br>(88.89, 98.22) | 97.26<br>(90.55, 99.51) | 88.89<br>(80.74, 93.82) | 96.55<br>(90.35, 99.06) | 97.26<br>(90.55, 99.51) |
| Wiener | 91.49<br>(84.10, 95.62) | 96.59<br>(90.45, 99.07) | 98.63<br>(92.64, 99.93) | 93.33<br>(88.84, 91.12) | 96.55<br>(90.35, 99.06) | 98.63<br>(92.64, 99.93) |
| InBios | 97.87<br>(92.57, 99.62) | 98.86<br>(93.84, 99.94) | 98.63<br>(92.64, 99.93) | 100.00<br>(95.91, 100.00) | 100.0<br>(95.77, 100.00) | 98.63<br>(92.64, 99.93) |

<sup>1</sup>Blood donors born in El Salvador (67), Guatemala (10), Honduras (7), Costa Rica (1), Nicaragua (1), unspecified Central America (2)
<sup>2</sup>Donors born in Bolivia (32), Argentina (13), Chile (5), Paraguay (2), Uruguay (1), Brazil (6), Colombia (9), Ecuador (2), unspecified South America (3)

**Figure 1.**
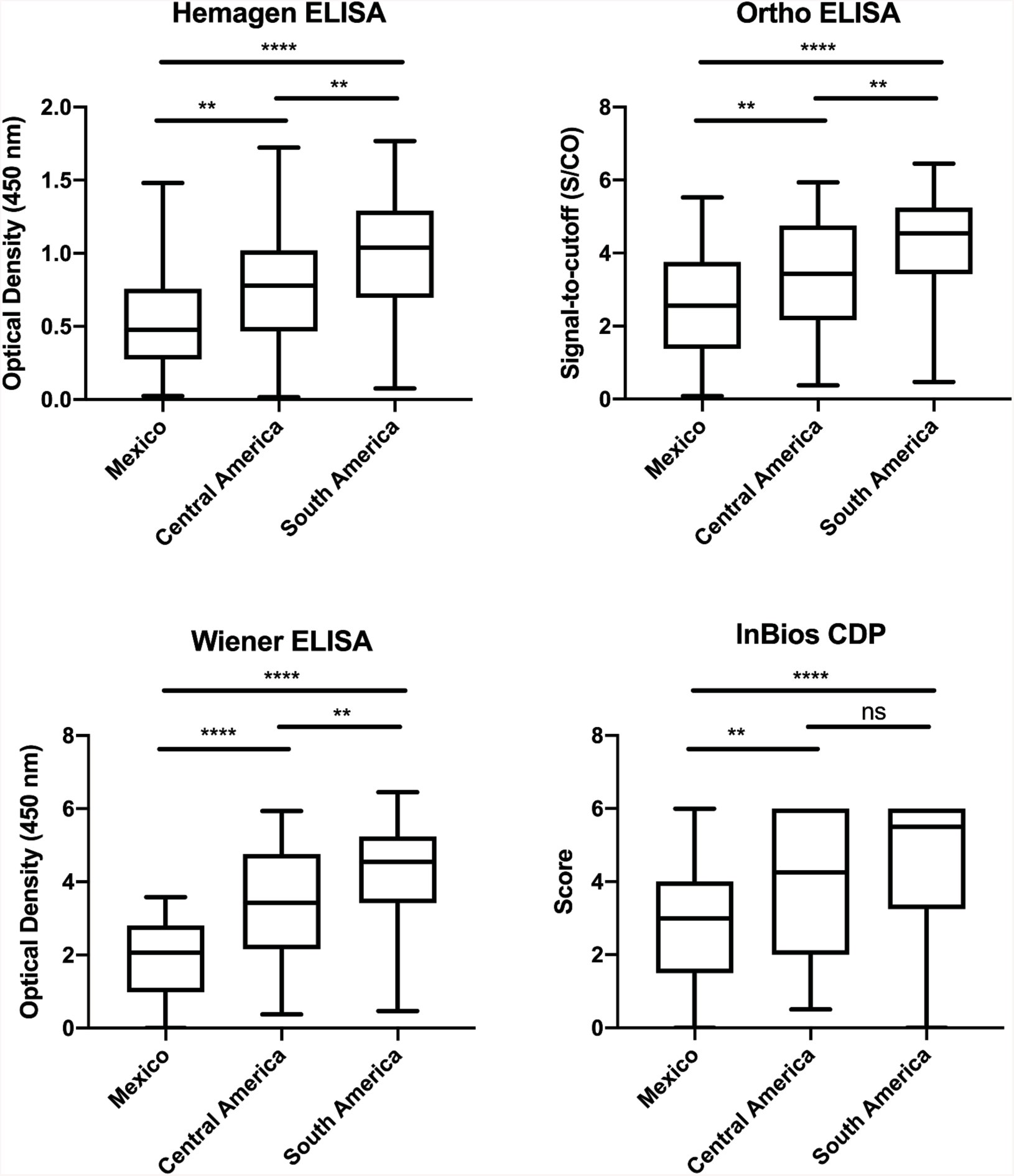
Distribution of positive serology values by blood donor region of birth. Results expressed as signal over cutoff (S/CO) for Ortho, optical density at 450 nm for Hemagen and Wiener, and score 0-6 for InBios. Across all tests, Mexican-born individuals showed the lowest test values and South American-born the highest. NS= P>0.05, *= P≤0.05, ** = P≤0.01, *** = P≤0.001, **** = P≤ 0.0001.

## DISCUSSION

Our data provide initial evidence for an appropriate diagnostic algorithm for Chagas disease in the US. The direct comparison of the four FDA-cleared tests demonstrates a range of sensitivity and specificity estimates across tests, and consistent variation in sensitivity by country of origin. Based on these findings, we can develop preliminary guidance for optimal use of these tests, anticipate associated challenges and identify where improvements are needed.

In common with recommendations for syphilis and early algorithms for HIV (25, 26), definitive diagnosis of chronic *T. cruzi* infection requires positive results by two distinct tests (3, 4). This algorithm was developed to address issues of both sensitivity and specificity. Simultaneous use of two tests optimizes both parameters and may be cost-effective in high prevalence settings. However, when low prevalence is anticipated, universal testing by two assays is impractical. Most programs will use one test as a screen and run only the screen-positives by the second assay. In these circumstances, the order is crucial; a high sensitivity screening test is essential to minimize the risk of missing true infections (Figure 2). At the same time, if specificity is not high, an assay will result in many false positives, potentially undermining confidence in testing. For example, in a setting of 1.5% prevalence (27), any specificity lower than 98.5% will result in more false than true positive results.

**Figure 2.**
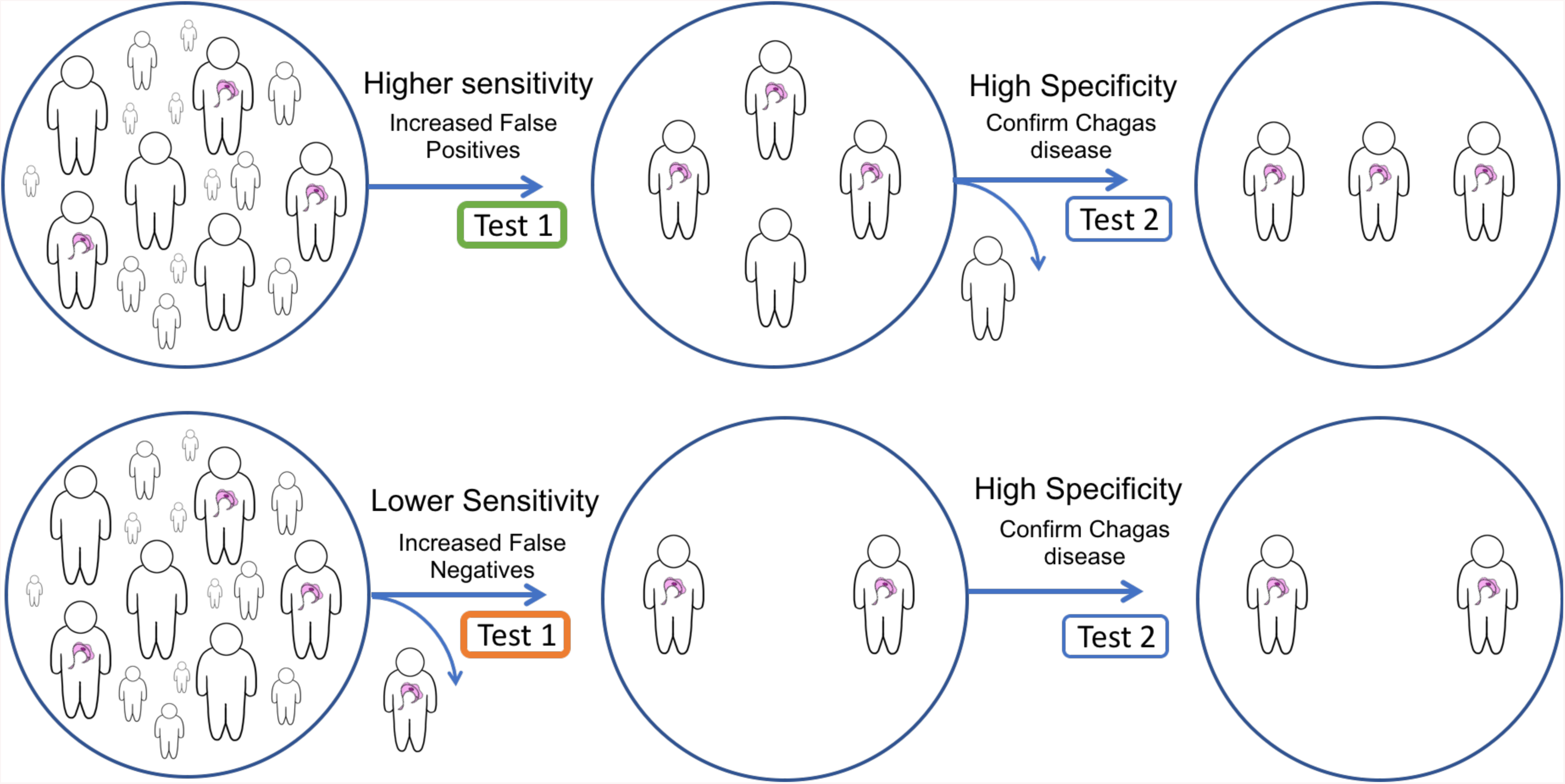
Effect of variation in clinical sensitivity of initial test in a two-step diagnostic algorithm. Two-step diagnostic algorithms allow for an acceptable number of false positives to ensure positive cases are detected. A) Illustrates higher sensitivity initial test, with a high specificity confirmatory test to rule out false positives. B) Illustrates a missed case of Chagas disease due to a lower sensitivity initial test and false negative result.

No single test had optimal performance characteristics in our data, despite the high sensitivity and specificity figures reported in their FDA 510(k) clearance applications and package inserts (28-31). In part, this may be attributable to the difference in performance in a setting closer to ‘real world’ diagnostic testing versus the more controlled setting of a clinical trial. However, a major issue in the available data is that few specimens from Mexico and Central America appear to have been included in preclinical testing (28, 29, 31). Only the Ortho evaluations reported results in specimens from Mexico, Guatemala and US at-risk populations during test development (19, 30). Published data confirm high rates of discordance and false-negative results by other assays in Mexico (16, 18) and the lower antibody reactivity seen in our data poses a challenge to achieving adequate sensitivity. Given the high proportion of US *T. cruzi* infections with Mexican origins, investigating and addressing the underlying cause of this phenomenon will be central to the effort to improve diagnostic test performance in the US. TcI, the predominant *T. cruzi* discrete typing unit (DTU) in Mexico, is widely distributed throughout the Americas (32). TcI also predominates in human infections in northern South America and Central America (33). Thus, the low reactivity in Mexican specimens is not a result of TcI predominance *per se*. Poorly understood strain differences within the TcI DTU may be responsible for the observed geographic variability in immune response (15, 17).

Based on the performance in our data, the Wiener Recombinante 3.0 and Ortho ELISAs showed the best balance of sensitivity and specificity, but both had suboptimal sensitivity in Mexican specimens. The InBios rapid test had the best sensitivity, with high sensitivity even in Mexican specimens, but its low specificity will result in a substantial number of false positives requiring confirmatory testing. The low sensitivity of Hemagen, especially in Mexican specimens, raises the risk of false negatives and concerns for its use as a screening test. In all cases, a discordant result between screening and confirmatory testing should prompt a third test as a tie-breaker, such as the IgG TESA-blot or the Abbott ESA, the latter having received FDA licensure for confirmatory use in the blood donor screening algorithm.

The use of surplus blood donation specimens has both limitations and advantages. Blood donor populations are not representative of the general US population; donors are younger and healthier than the population at large, and although the rate of donation by Hispanics has increased markedly over the past decade, this group remains underrepresented (34, 35). However, given the design of the study, these differences should not affect the validity of the test performance estimates. Although three of the four tests are validated for both serum and plasma, the Hemagen package insert specifies the use of serum; we had only plasma available, which may have had an impact on our estimates for this assay. However, other IgG ELISA tests have reported equivalent results in plasma and serum (36). The decrease in reactivity by the Ortho ELISA in current vs prior BD testing is perplexing. Length of storage was inversely related to the magnitude of the decline, making antibody degradation an unlikely explanation. The Ortho ELISA uses cultured parasite lysate as its antigen source, possibly introducing biological variability.

A critical review of diagnostic studies suggests that a double-blinded prospective cohort provides the optimal study design, because testing of positive and negative groups selected on the basis of prior test results introduces a bias toward overestimates of performance characteristics, especially if discordant specimens are excluded (37). However, prospective testing by multiple assays in a very low prevalence population would incur prohibitive costs. We attempted to minimize bias by including specimens that were discordant in BD testing and specimens across the entire range of antibody responses, and by using two different comparators (BD and consensus) and a latent class analysis. Our results demonstrate how performance estimates may vary, depending on the comparator and analysis method. Our design was strengthened by the large sample size, robust set of low titer positives and infections acquired in different geographic regions, characteristics difficult to replicate in the US in the absence of a large, well-funded multicenter study. Our results do not preclude such a study. On the contrary, additional rigorous analyses of data from robust specimen sets with broad geographic coverage are essential to better understand and improve the performance of the available tests in US populations at risk of *T. cruzi* infection.

## CONCLUSION

In an analysis of US blood donor specimens, InBios Chagas Detect Plus rapid test had the highest sensitivity but lowest specificity, while Hemagen had the lowest sensitivity of the FDA-cleared tests. Hemagen, Ortho, and Wiener ELISAs all had equivalently high specificity. Sensitivity was lower for the Ortho, Wiener and Hemagen ELISAs in specimens from donors born in Mexico, intermediate for Central America and highest for South America, consistent with differences in the distribution of antibody reactivity in these groups. Use of a high sensitivity screening test, followed by a second higher specificity test, offers the best current algorithm for diagnostic screening in the US.

## Acknowledgements

We thank Dr. Yagahira Elizabeth Castro Sesquen for sharing her semi-quantitative scale for scoring InBios test results, InBios International and Wiener Laboratories for donation of test kits, and Professor Charles McCullough for his advice on the use of latent class analysis.

Potential conflicts of interest. CB reports consulting fees from Exeltis in 2018. All other authors (JW, CAB, EG, AI, RT, SS, JS) reported no conflicts of interest.

## Funding

This study was supported by the Mundo Sano Foundation. CB receives partial salary support from Mundo Sano Foundation. The participation of JW was supported in part by a grant from the National Heart, Lung, and Blood Institute at the National Institutes of Health under award number R38 HL143581. The participation of CAB, EG, and JS was supported in part by the Bill and Melinda Gates Foundation under award number OPP1017584. The funding sources had no role in the study design, collection, analysis and interpretation of the data, preparation of the manuscript, or the decision to submit for publication. This publication’s contents are solely the responsibility of the authors and do not necessarily represent the official views of their sponsors.

